# Nanopore metagenomic sequencing links clinically relevant resistance determinants to pathogens

**DOI:** 10.64898/2026.02.16.706128

**Authors:** Harika Ürel, Ela Sauerborn, Michael Biggel, Friedemann Gebhardt, Ebenezer Foster-Nyarko, Silvio D Brugger, Rhys T White, Søren Heidelbach, Mads Albertsen, Francis Muchaamba, Tim Reska, Marc J A Stevens, Roger Stephan, Richard Fetherston, Lara Urban

## Abstract

Culture-independent metagenomics enables the detection of plasmid-encoded antimicrobial resistance (AMR) genes directly from clinical samples; however, the clinical significance of these genes depends on their bacterial host and genomic context, which metagenomics cannot fully infer. Nanopore sequencing technology intrinsically encodes epigenetic modifications such as methylation, which can be leveraged for plasmid-host associations from metagenomic data. Existing methods rely on the recovery of metagenome-assembled genomes (MAGs), which can introduce bias toward abundant taxa and leave clinically relevant, low-abundance pathogens unassociated. To address this limitation, we extended methylation-based plasmid-host association from the MAG level to individual assembly contigs and sequencing reads. The CUPID pipeline implements the calculation of contig and read similarity scores, which compare weighted mean methylation rates across motifs genetically shared between any contig or read pair. We validated this approach on a mock metagenomic community composed of ten carbapenem-resistant *Enterobacterales* isolates, where we achieved 93.8% accuracy at the contig level and 100% at the read level for carbapenemase plasmid-host associations. When applied to metagenomic and quasimetagenomic data of sixteen patient rectal swabs collected during routine hospital surveillance, our approach assigned every detected plasmid-encoded carbapenemase to its correct bacterial host at the contig level, using matched culture-based diagnostics and whole-genome sequencing as a ground truth. Read-level analysis identified additional associations that were missed at the contig level, including a multi-host plasmid confirmed by established diagnostics. These findings demonstrate a pathway from rapid AMR gene detection using metagenomics to actionable surveillance for infection prevention, transmission tracing, and outbreak investigation.

**Impact statement:** Culture-independent metagenomics can detect antimicrobial resistance genes, but their clinical significance depends on the bacterial host and genomic context. Here, we show that nanopore-derived bacterial DNA methylation patterns can link carbapenemase genes to pathogenic hosts and plasmid context directly from patient samples. This provides a route from rapid antimicrobial resistance gene detection to actionable public health surveillance.

**Data summary:** All sequencing data after human content filtering have been deposited at the European Nucleotide Archive (ENA, BioProject accession PRJEB108076, with all isolate sequencing data for mock community generation available under the sample accession numbers SAMEA121375149-58, all isolate sequencing data from the rectal swabs available at SAMEA121334008-24, all metagenomic data from the rectal swabs available at SAMEA121325220-27, and all quasimetagenomic data available at SAMEA122914816-23, SAMEA122920068-74). All code is available at GitHub: https://github.com/harikaurel/cupid. All other supporting data are provided in the article and supplementary tables.

## Introduction

Genomics is now central to infectious disease surveillance, supporting high-resolution outbreak detection, transmission inference, source attribution, and antimicrobial resistance (AMR) monitoring across public health and food safety systems (1,2). Whole-genome sequencing (WGS) of cultured pathogens is now, for example, a recommended standard by the European Centre for Disease Prevention and Control and the European Food Safety Authority (3,4). Portable, real-time nanopore sequencing is extending these capabilities beyond centralized sequencing centers, making pathogen genomic surveillance increasingly feasible in clinical microbiology laboratories, distributed public health surveillance networks, and lower-resource settings (5–9).

In comparison to culture-based genomic characterization, rapid, culture-independent metagenomic pathogen surveillance offers several advantages. Metagenomic sequencing can be applied to a wide range of clinical sample types without culture-based enrichment, which can be time-consuming, requires specialized biosafety infrastructure, and depends on pathogen-specific growth requirements (3). Metagenomics can detect pathogens and their subtypes, including the identification of unculturable or previously uncharacterized pathogens through *de novo* genome assembly approaches. Metagenomic data can further detect virulence- and AMR-associated genes and, where sequence linkage permits, place them in pathogen or mobile-element contexts (such as plasmids), including those harbored within commensal microbial communities (10,11). Recent advances in accurate long-read metagenomics using nanopore sequencing technology can provide (*i*) higher resolution for read-based taxonomic classification and AMR annotations, (*ii*) more contiguous *de novo* assemblies (contigs) from complex microbial environments, including the recovery of circularized AMR-carrying plasmid contigs, and (*iii*) binning of these contigs into high-quality metagenome-assembled genomes (MAGs), enabling the reconstruction of near-complete or closed genomes by binning of the recovered contigs (5,12).

While nanopore metagenomic sequencing has recently demonstrated improved pathogen and AMR profiling compared with culture-based approaches (10), three analytical barriers continue to limit the clinical interpretation of metagenomic data: (*i*) insufficient sensitivity for detecting low-abundance pathogens and AMR genes; (*ii*) difficulty distinguishing clinically relevant organisms from background or non-viable DNA; and (*iii*) limited ability to link detected AMR genes to their bacterial hosts, particularly when genes are plasmid-borne. The properties of nanopore sequencing data may provide opportunities to overcome these barriers: Nanopore sequencing measures disruptions in an ionic current as native DNA molecules pass through a nanopore, generating raw signal data that can be basecalled into nucleotide sequence while retaining information about molecule-level features such as epigenetic modifications (13). (*i*) To increase sensitivity, the real-time signal acquisition of nanopore sequencing enables adaptive sampling, in which the initial sequence from each molecule is rapidly basecalled and aligned during sequencing so that only molecules matching predefined pathogen or AMR targets are retained (14–16). (*ii*) The raw nanopore signals from native DNA sequencing can support additional computational inference beyond consensus sequence alone. Ürel et al. (2025) showed that signal-level features can be used to infer microbial viability in controlled experimental and mock-community settings, suggesting a possible route to distinguishing DNA from viable and non-viable organisms in metagenomic samples (17). (*iii*) Finally, native nanopore sequencing can detect bacterial DNA methylation patterns, which can be used as endogenous genomic signatures for metagenomic binning and plasmid-host associations (18). This approach leverages bacterial restriction-modification systems, in which methyltransferases methylate specific host DNA motifs while cognate restriction enzymes cleave unmethylated foreign DNA (19, 20). The Nanomotif tool exploits the methylation motifs using nanopore sequencing data to identify methylated sequence patterns within contigs, improve metagenomic binning, and associate unbinned contigs, including plasmids, with host MAGs (18, 21, 22).

Linking plasmids to their pathogenic hosts is critical when detecting plasmid-encoded AMR genes in metagenomic data. Carbapenemase-producing *Enterobacterales* exemplify plasmid-mediated AMR spread, with conjugative and mobilizable plasmids moving KPC, NDM, VIM, and OXA-48-like genes across bacterial strains and species (23). Given that delays in diagnosis and therapy of carbapenemase-producing *Enterobacterales* infections are associated with high morbidity and mortality, rapid metagenomic identification of pathogens, AMR determinants, and their genomic context could substantially improve infection management and public health response (24, 25). However, existing methylation-based plasmid-host association approaches depend on the recovery of the prospective microbial host on the MAG level (18), which is poorly matched to clinical surveillance samples where the microbial hosts of AMR determinant may be present at low abundance within a complex mixture of host DNA, dominant commensal microbiota, and other microbial DNA. Applying Nanomotif directly to carbapenemase-carrying plasmids in low-abundance clinical metagenomes is therefore constrained, with its plasmid-host association workflow depending on sufficient recovery of the prospective host MAG, while pathogens are mostly only detectable at the contig or sequencing read level. Consequently, the use of nanopore-based methylation information in metagenomics has so far provided limited evidence of clinical utility and has not yet been validated against established culture-based diagnostics (18,21,22).

Here, we developed and validated a computational framework that extends methylation-based plasmid-microbial host associations to the assembly and individual sequencing read level. We used mock metagenomic data from clinically relevant carbapenem-resistant *Enterobacterales* isolates to validate associations between plasmid and chromosomal contigs and reads predicted by our computational framework. We then evaluated this framework on metagenomic data derived from rectal swabs routinely collected from patients during clinical surveillance for carbapenem-resistant *Enterobacterales* carriage. By comparing our predictions against pathogen and carbapenemase predictions from established culture-based diagnostics and matched whole-genome sequences, we show that nanopore-based metagenomics can reliably distinguish plasmid-encoded AMR genes and associate them with their respective bacterial hosts, increasing the sensitivity and precision of metagenomics-based pathogen surveillance.

## Methods

### Mock metagenomic community

We generated mock metagenomic communities by pooling the WGS data from ten carbapenem-resistant *Enterobacterales* isolates. These mock communities were used to benchmark metagenomic associations between AMR genes and plasmids, with the matched whole genomes as ground truth.

#### DNA extraction

Ten carbapenem-resistant *Enterobacterales* isolates (Table S1A) were recovered from glycerol stocks stored at -80 °C and cultured overnight at 37 °C on BD® Columbia Blood Agar plates. Genomic DNA was extracted from the isolates using a spin-column-based purification method. Briefly, 10–20 colony-forming units (CFUs) from overnight cultures were resuspended in 1 mL phosphate-buffered saline (PBS). A 100 µL aliquot was adjusted to 1 mL with PBS and centrifuged at 12,000 × g for 2 min. Genomic DNA was extracted from the resulting pellet using the DNeasy Blood & Tissue Kit (QIAGEN, Hilden, Germany), according to the manufacturer’s protocol for Gram-negative bacteria (26). To improve DNA yield and purity, 40 µL of proteinase K was added, and the lysis incubation was extended to 2 h. An RNase treatment step was incorporated after proteinase K incubation, following the manufacturer’s instructions (Qiagen, Hilden, Germany) (8). DNA concentrations were measured using a Qubit 4 Fluorometer (dsDNA BR kit).

#### Whole-genome generation by nanopore sequencing

Nanopore sequencing libraries were prepared using the Oxford Nanopore Rapid Barcoding Kit (SQK-RBK114.96). Two barcodes were used per isolate to adjust for imbalanced sequencing across barcodes. Per barcode, 200 ng of DNA was used. The final libraries were sequenced on R10.4.1 PromethION flow cells for 72 h. The nanopore sequencing raw data were basecalled using the Dorado v0.9.1 super-accuracy basecalling model v5.0.0 (dna_r10.4.1_e8.2_400bps_sup@v5.0.0) and with epigenetic modification calling enabled for 6mA, 4mC, and 5mC. Reads were demultiplexed using Dorado demux and filtered to retain reads with a minimum Phred quality score of 9 and a minimum length of 200 bases using Chopper v01.11.0 (28). Sequencing summary metrics were generated using SeqKit v2.10.1 (29). *De novo* genome assemblies were generated using Flye nano-hq v2.9.6 (30, 31). Assemblies were subsequently polished using Dorado polish in bacterial mode. Assembly depth was assessed using SAMtools depth v1.21 (32). Genome quality metrics, including assembly completeness and contamination, were evaluated using CheckM2 v1.1.0 with the *uniref100.KO.1.dmnd* reference database (33). Polished *de novo* assemblies were analyzed using the Pathogenwatch v2.3.1 platform (accessed 28 January 2026) for species identification and, if available, multi-locus sequence typing (MLST) (34). Assemblies were further taxonomically classified using Kraken2 v2.1.3 (35) and the NCBI nucleotide database (accessed 30 January 2026). AMR genes were identified at the contig level using AMRFinderPlus v4.0.23 (36–38) (database version 2024-10-22.1). Contigs were annotated as plasmids or chromosomes using MOB-suite (v3.1.9) (39), which also provides plasmid typing and mobility predictions based on replicon markers, relaxase (MOB) typing, and mating-pair formation (MPF) systems. The MOB-recon module was applied specifically to AMR-carrying contigs classified as plasmids to determine plasmid incompatibility types.

### Clinical metagenomics from rectal swabs

Sixteen rectal swabs were routinely collected as part of the Technical University of Munich hospital admission screening for carbapenem-resistant *Enterobacterales*. Swabs (Copan Diagnostics, Brescia, Italy) were inserted rectally to a depth of 2–3 cm and rotated three times. Following collection, swabs were plated onto BD® MacConkey agar (Becton Dickinson GmbH, Heidelberg, Germany) and Thermo Scientific™ Brilliance™ extended-spectrum β-lactamase agar plates and incubated at 37 °C for 20–24 h.

#### Established diagnostics

Species identification was performed from a single CFU of *Enterobacterales* isolates per rectal swab (Tables S1B&D) using matrix-assisted laser desorption ionization time-of-flight mass spectrometry (MALDI-TOF MS; Bruker Daltonics GmbH, Bremen, Germany; MBT Compass 4.1 with MBT Compass Reference Library 2023), according to the manufacturer’s instructions. Phenotypic antimicrobial susceptibility testing (AST) was conducted on pure subcultures using the VITEK® 2 system (bioMérieux, Marcy l’Etoile, France) with the AST-GN69 cards, and carbapenem susceptibility was interpreted according to European Committee on Antimicrobial Susceptibility Testing (EUCAST) guidelines (27). Vitek® 2 reports categorical susceptibility interpretations [susceptible/intermediate/resistant (S/I/R)] derived from discrete antimicrobial concentration steps; where we refer to elevated non-susceptibility, we mean a shift from S to I/R in the Vitek^®^ 2 categorical call. Carbapenemase production was assessed using a multiplex immunochromatographic lateral flow assay (O.K.N.V.I Resist-5) that detects the presence of the most common KPC, OXA-48-like, NDM, VIM, and IMP carbapenemases. After completion of routine diagnostics, five to ten CFUs of each carbapenem-resistant *Enterobacterales* isolate were stored at -80 °C in glycerol stocks. The same isolates were thawed and processed for WGS as described above.

#### Metagenomic data generation

Genomic DNA was extracted directly from the first eight (of overall 16) collected rectal swabs (“metagenomic” samples), and from a negative extraction control (PBS). These swabs, stored in Amies transport medium without charcoal (Copan Diagnostics, Brescia, Italy), were thawed at 4 °C overnight. For lysis, the swab tip was transferred into a QIAamp PowerFecal Pro DNA Kit PowerBead tube. The swab shaft was aseptically removed using sterile scissors and tweezers, ensuring that no residual transport medium or gel was transferred. The PowerBead tube with the swab tip was vortexed at high speed for 1 min. Subsequently, the QIAamp PowerFecal Pro DNA Kit solution CD1 was added according to the manufacturer’s instructions, followed by incubation at 65 °C for 10 min and bead beating. DNA extraction was then completed according to the QIAamp PowerFecal Pro DNA Kit protocol. DNA concentrations were measured using the Qubit 4 Fluorometer (dsDNA BR kit). Nanopore sequencing libraries were prepared with the SQK-RBK114.24 Rapid Barcoding Kit and sequenced for 72 h on a MinION Mk1D device using two R10.4.1 MinION flow cells each. Two barcodes were used per sample per DNA extract. A total of 40-200 ng of DNA in 10 µL per barcode was used for each library preparation, depending on the respective DNA concentration.

The remaining eight rectal swabs were incubated in 4 mL of brain heart infusion (BHI) broth for 4 h at 36 °C (“quasimetagenomic” samples). Following incubation, a 1 mL aliquot of the suspension was collected and centrifuged at 12,000 × g for 5 minutes. The resulting pellet was resuspended in 100 µL of the remaining BHI. DNA extraction and nanopore sequencing were performed as previously described for the metagenomic samples.

### Metagenomic data analysis

The entire computational pipeline is available at GitHub: https://github.com/harikaurel/cupid, including the Nanomotif-based CUPID pipeline for the implementation of *contig similarity score* (css) and *read similarity score* (rss) calculations and visualizations.

#### Read-level metagenomic data processing

All nanopore sequencing data were basecalled using Dorado v0.9.1 with the SUP v5.0.0 basecalling model (dna_r10.4.1_e8.2_400bps_sup@v5.0.0) and with epigenetic modification calling enabled for 6mA, 4mC, and 5mC. Reads were demultiplexed using Dorado demux and filtered to retain reads with a minimum quality score of 9 and a minimum length of 200 bases using Chopper v01.11.0 (28). Sequencing summary metrics were generated using SeqKit v2.10.1 (29). To remove human genomic information, human sequencing reads were identified by mapping to the human reference genome (GRCh38) using minimap2 v2.30 (40) with the map-ont preset. Reads with human alignments were excluded, and unmapped reads were retained as the host-depleted metagenomic dataset. Read-level AMR gene detection was done using AMRFinderPlus v4.0.23 (36). Plasmid reads were detected by MOB-suite (v3.1.9) (39), and read-level taxonomic classification was performed using Kraken2 v2.1.3 (35) and the NCBI nucleotide database (accessed 30 January 2026).

#### Metagenomic assemblies and binning

For metagenomic assemblies (Figure 1A), we used nanoMDBG (v.1.0) (42,43) followed by polishing using Dorado aligner and polish (in bacterial mode). Contig-based taxonomic classification was performed with Kraken2 as described above. AMR genes were identified using AMRFinderPlus v4.0.23 (36) as described above. In the mock metagenomic community, the ground truth for each metagenomic contig was determined by aligning the sequencing reads from each isolate to the mock community assemblies with minimap2 v2.30 (map-ont preset) (40), retaining only primary alignments.

**Figure 1.**
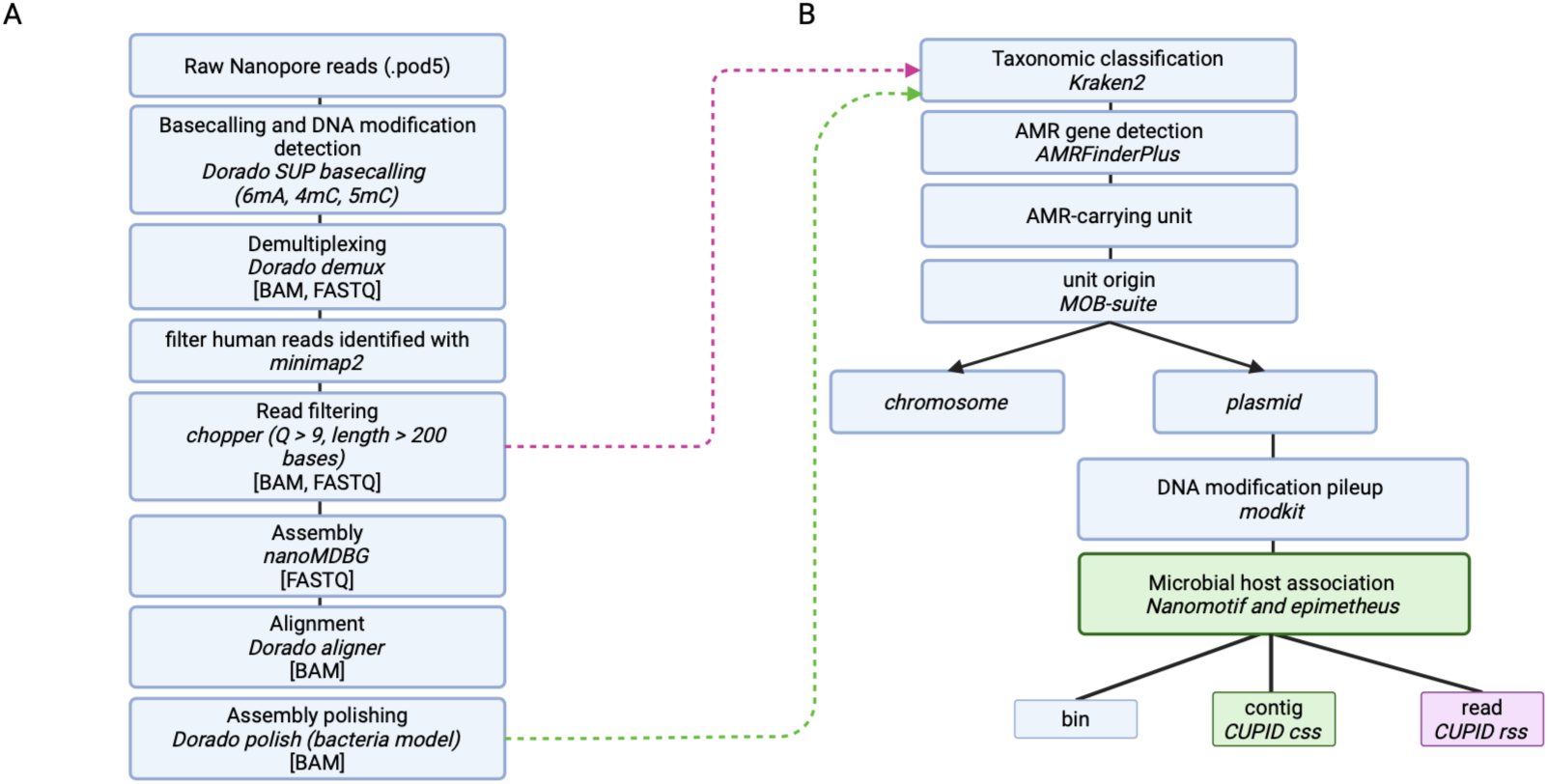
Computational pipeline for nanopore metagenomic data analysis and plasmid-microbial host associations. **A.** Nanopore metagenomic data analysis: Raw sequencing data are basecalled using Dorado super-accuracy basecalling (SUP), incorporating the detection of epigenetic modifications (6mA, 4mC, 5mC). Reads are demultiplexed, filtered for human origin, quality (Q), and length, and assembled using nanoMDBG. Assemblies are polished into final metagenomic contigs. **B.** Association analysis of individual sequencing reads or contigs (“units”): Taxonomic classification is performed using Kraken2 and origin (chromosomal or plasmid) of all AMR-carrying units is assigned using MOB-suite. For chromosomally annotated AMR genes, host associations are inferred through taxonomic classification of the respective contig or read. For AMR-carrying plasmid units, their methylation information (modkit) is input into Nanomotif and epimetheus tools; while standard Nanomotif allows for bin-level associations, the here-implemented CUPID pipeline allows for contig- and read-level associations using contig and read similarity scores (css and rss).

Plasmid replicon types and mobility features were inferred from contigs using MOB-suite (v3.1.9) (Figure 1A). These plasmid contigs were additionally manually investigated to rule out false-positive plasmid annotations: (*i*) for the presence of genes involved in plasmid and/or transposon function using Prokka (43), and (*ii*) for the occurrence of the sequence in at least two strains after BLAST alignment against the full NCBI database at ≥95% nucleotide identity (44). Finally, the plasmids were annotated for their insertion sequence (IS) elements by BLAST-based homology search against the ISfinder nucleotide database (downloaded on May 26th, 2026), requiring ≥80% nucleotide identity (45). IS detections were deduplicated by the IS family to reduce redundancy arising from the high sequence similarity between IS family members, where detections from the same IS family within 50 bases of each other were considered to represent a single insertion event. Each carbapenemase detection was then evaluated for the presence of IS elements within a conservative 500 bases-window upstream (5’) and downstream (3’) of the gene boundaries, and putative composite transposons were identified where IS elements of the same family were detected in inverted orientation on both sides of the respective gene.

Contig binning was performed using MaxBin2 (v2.2.7; 46), MetaBAT2 (v2.18; https://bitbucket.org/berkeleylab/metabat), and VAMB (v5.0.4; *vamb bin* command with *–m* 2000 and *–minfasta* 200000; https://github.com/RasmussenLab/vamb). The resulting bins were integrated using DASTool (v1.1.7; https://github.com/cmks/DAS_Tool).

#### Plasmid-microbial host associations

Per-base epigenetic modification counts per sample were generated in BED format using the modkit pileup function (v0.5.0; https://github.com/nanoporetech/modkit). We then used this BED file and the FASTA file of the metagenomic assembly per sample to apply the Nanomotif v0.8.0 (18) motif_discovery module and identify methylation motifs at the contig level. As Nanomotif is normally applied to the bin level, we implemented the Nanomotif-based CUPID pipeline for contig- and sequencing read-level association calculations (https://github.com/harikaurel/cupid).

For contig-level associations (Figure 1B), we first modified the contig-to-bin assignment file such that each contig was treated as its own bin (i.e., one contig per bin). Chromosomal contigs that remained “Unclassified” by Kraken2 were discarded. We adjusted several default Nanomotif parameters to make the motif detection more lenient and recover more putative motifs from the metagenomic data (setting min_motifs_bin to 1, threshold_valid_coverage to 1, and --min_motif_score to 0.5). Motifs of a length of at least four bases in the resulting bin-motifs.tsv are used as input for the epimetheus tool (epimetheus methylation-pattern contig function), which generates the motifs-scored-read-methylation.tsv output containing weighted mean methylation scores per motif and contig. We used these methylation scores to calculate pairwise contig distances using root mean square distance (*RMSD*), which is calculated based on all genetically shared motifs between any AMR-carrying plasmid and chromosomal contig. We then calculated the *contig similarity score* (css) as *RMSS x n*, where *RMSS* is calculated as (1 − *RMSD*) and *n* denotes the number of shared motifs between any contig pair. While the taxonomic annotation of the chromosomal contig with the highest css is chosen as putative host of the respective plasmid contig, CUPID additionally visualizes the overall contig similarity structure to understand the robustness of these plasmid-host associations. To do so, CUPID also calculates the css between all chromosomal contigs and then visualizes the contig similarity structure with reference to the AMR-carrying plasmid contig using Principal Coordinate Analysis (PCoA; using python scikit-bio v0.5.9). It additionally generates heatmaps of per-motif weighted mean methylation values, comparing the plasmid against the top-scoring chromosomal contig of each of the top five species-level taxa according to the css. The number in parentheses in the heatmap cells indicates the number of motif observations per contig.

Read-level associations follow the same principle as contig-level associations to calculate the *read similarity score* (rss), with the following exceptions: The motifs identified at the contig level are used and supplied to the epimetheus methylation-pattern read-bam module, which reports the per-read methylation quality of each motif at each position in the read as extracted from the ML tag of the BAM file. These quality values are converted to probabilities by dividing by 256, giving a range of [0, 1]. For each read, the mean methylation probability across the positions of a given motif is reported as the read’s mean methylation value.

## Results

### Standard clinical metagenomic analyses

Metagenomic data was generated from 16 rectal swabs, which were routinely collected as part of the Technical University of Munich hospital admission screening for carbapenem-resistant *Enterobacterales* (Methods). The presence of carbapenem-resistant *Enterobacterales* in the samples was confirmed using established culture-based diagnostics and WGS (Table S1A; Methods); half of the swabs (n=8) were processed directly (“metagenomic” samples) and the other half of the swabs (n=8) were processed after short unselective enrichment (“quasimetagenomic” samples) to increase metagenomic sensitivity (Table S1B; Methods). We restricted our metagenomic AMR gene-host association analysis to carbapenemases and their respective *Enterobacterales* hosts since these associations could be verified using established diagnostics and WGS data of the respective isolates (Methods).

When applying traditional metagenomic assembly and binning to all (quasi)metagenomic data, we found that binning recovered a pure MAG of the relevant carbapenem-resistant *Enterobacterales* species in only three of the 16 samples, with one of those three relevant MAGs only consisting of a single contig (Methods; Table S2). We therefore moved the inference of plasmid-host associations from the MAG to the contig and individual sequencing read level by developing a *contig similarity score* (css) and *read similarity score* (rss) that compare weighted mean methylation rates and shared motif profiles between individual contigs or sequencing reads instead of on the MAG level (Methods). The calculation of these scores is implemented in the CUPID pipeline (Methods).

### Contig- and read-level plasmid-microbial host associations

#### Application to mock metagenomic community

We generated mock metagenomic data to first validate predicted plasmid-microbial host associations. This data comprised pooled WGS data from ten clinically relevant carbapenem-resistant *Enterobacterale*s isolates, each with accompanying near-complete chromosomal and plasmid assemblies (Table S1C; Methods). We assembled the mock community (Table S1C), calculated the css and rss of all plasmid units carrying NDM, KPC, OXA-48, or VIM carbapenemases (Figure 1), and compared the predicted microbial host associations with the WGS-based ground truth (Table 1; Methods). This approach achieved high accuracy of 93.8% (15/16) in plasmid-host association predictions on the contig level; the only AMR-carrying plasmid contig that could not be associated with its correct microbial host was a hybrid assembly (Table S3A). At the read-level, all carbapenemase gene variants and families were linked to their expected microbial host (100%; 8/8). For NDM, rss resolved both expected hosts in the mock community, assigning 78 NDM-carrying reads to *Proteus mirabilis* and 54 reads to *E. coli* (Table S4A).

**Table 1.** Contig- and read-level associations between carbapenemase-carrying plasmid contigs and their putative microbial host according to *contig* and *read similarity scores* (css and rss) across eight metagenomic (M1-8) and eight quasimetagenomic (Q1-8) datasets (C=Carbapenemase; P=Pathogen host). For contig-level associations, the contig-encoded carbapenemase gene variant or family and gene location and the taxonomic annotation of the chromosomal contig with the highest css are indicated. For read-level associations, each carbapenemase-carrying read is assigned to the host read with the highest rss; taxonomic annotations are shown per gene variant or gene family (number of supporting associations in parentheses; Table S4B). The established diagnostics (“Diagnostics”) and whole-genome sequencing (“WGS”) results of all carbapenem-resistant *Enterobacterales* grown from the respective samples are included as ground truth. The carbapenemases were determined by lateral flow test and the pathogenic host by MALDI-TOF MS (“c.”= complex); where several carbapenemases and/or several pathogenic species are indicated in one cell, all carbapenemases were detected in all of the respective isolates. The WGS results include the multi-locus sequence type (ST) per pathogenic species in parentheses. (*OXA gene copies were detected on the contig and read level and associated with *Citrobacter freundii*, but they were assigned to the OXA-1-family in follow-up analyses. **Ambiguous association at read level. ***Taxonomic annotation of the contig/read that carries the chromosomally encoded OXA-244 gene, i.e., no methylation-based association. ****Only metagenomics-based association not recovered by the isolate ground truth (Sample Q4, KPC-2, read-level). *****No gene variant assignment possible due to detection at the contig edge (90.9% reference coverage).

| # | Metagenomics |  |  |  | Isolate ground truth |  |  |  |
| --- | --- | --- | --- | --- | --- | --- | --- | --- |
|  | Contig-level |  | Read-level |  | Diagnostics |  | WGS |  |
|  | C | P | C: P (# reads) |  | C | P | C | P |
| M1 | N.A.* |  |  |  | OXA-48-like | C.freundii c. | OXA-48 (IncL/M) | C.freundii (ST22) |
| M2 | N.A. |  |  |  | KPC | C.freundii c. | KPC-2 (IncN) | C.freundii (ST256) |
| M3 | OXA-48 (IncL/M) | C.freundii | OXA-48 | C.freundii (1)** | OXA-48-like | C.freundii c. | OXA-48 (IncL/M) | C.freundii (ST22) |
| M4 | N.A. |  |  |  | KPC | K.variicola | KPC-2 (IncN) | K.variicola (ST347) |
| M5 | KPC (plasmid) | K.pneumoniae | KPC | E.coli (1) | KPC | K.pneumoniae, E.coli | KPC-2 (IncN) | K.pneumoniae (ST147), E.coli (ST*5a6) |
| M6 | OXA-244 (chrom) | E.coli*** | OXA-244 | E.coli*** | OXA-48-like | E.coli | OXA-244 (chrom) | E.coli (ST 58/87) |
| M7 | OXA-244 (chrom) | E.coli*** | OXA-244 | E.coli*** | OXA-48-like | E.coli | OXA-244 (chrom) | E.coli (ST69/3) |
| M8 | N.A. |  |  |  | NDM | K.pneumoniae | NDM-5 (IncFIB) | K.pneumoniae (ST395) |
| Q1 | VIM-1 (IncL/M) | E.coli | VIM: K.pneumoniae (4)<br>Oxa-48: E.coli (3)<br>Oxa-48-like: K.pneumoniae (2) |  | VIM<br>Oxa-48-like | E.cloacae c., K.pneumoniae, E. coli | N.A. |  |
| Q2 | KPC-2 (IncFII, IncN) | K.pneumoniae | KPC-2: K.pneumoniae (27)<br>KPC: K.pneumoniae (98) |  | KPC | K.pneumoniae | KPC-2 (IncN) | K.pneumoniae (ST147) |
| Q3 | NDM-1 (IncN) | E. coli | NDM-1: E. coli (15)<br>NDM: E. coli (74) |  | NDM | E.cloacae c., E. coli | NDM-1 (IncN) | E.coli (ST*1c08) |
| Q4 | KPC-2 (IncN) | K. pneumoniae | KPC-2: E.coli (1)****<br>KPC: K.pneumoniae (2) |  | KPC | K.pneumoniae | KPC-2 (IncN) | K.pneumoniae (ST147) |
| Q5 | OXA-48 (IncL/M) | C. freundii | OXA-48: C.freundii (3)<br>OXA-48-like: C.freundii (10) |  | OXA-48-like | C.freundii c. | OXA-48 (IncL/M) | C.freundii (not typable) |
| Q6 | KPC-2 (IncN) | E. hormaechei | KPC-2: E.hormaechei (25)<br>KPC: E.hormaechei (79) |  | KPC | E.cloacae c. | KPC-2 (IncN) | E.hormaechei (ST 102) |
| Q7 | KPC-2 (IncN) | E. coli | KPC: E.colii (4) |  | KPC | E.coli | KPC-2 (IncN) | E.coli (ST2541) |
| Q8 | NDM-5 (IncFIB, IncHI1B), OXA-48-like***** (IncL/M) | K.pneumoniae | NDM-5: K.pneumoniae (3)<br>NDM: K.pneumoniae (14)<br>OXA-48: K.pneumoniae (16) |  | OXA-48-like<br>NDM | K.pneumoniae, E. coli | NDM-5 (IncFIB, IncHI1B)<br>OXA-48 (IncL/M) | K.pneumoniae (ST383) |

#### Application to clinical metagenomics—Contig-level associations

We then assessed the contig- and read-based plasmid associations in the rectal swab (quasi)metagenomic data (Table S1B; Methods). We detected the relevant carbapenemases on the contig and read level in four (M3,5,6,7) of the eight (50%) metagenomic samples, and in all (Q1-8) of the eight (100%) quasimetagenomic samples (Table 1; see Table S3B for all other AMR genes).

Of the overall 13 carbapenemases (Q8 encoded two carbapenemases), two were predicted to be chromosomally encoded (M6,7); these chromosomally encoded *bla*^OXA-244^ genes were associated with *E.coli* according to the taxonomic classification of the gene-carrying contig itself, which could be confirmed by WGS of the respective *Enterobacterales* isolate (Table 1). The remaining 11 carbapenemases were predicted to be encoded by plasmid contigs; all of them could be associated with their correct microbial host according to isolate-based established diagnostics and WGS using our css (Table 1). In the quasimetagenomic data, every sample recovered a carbapenemase-carrying plasmid contig that could be correctly associated with its host using the css. The resolution of the metagenomic contigs was on par with the WGS assemblies and surpassed the resolution of established diagnostics such as MALDI-TOF MS and lateral flow tests: The metagenomic assemblies correctly classified pathogens to the species level and the carbapenemases to the gene variant level, identified the correct carbapenemase location (plasmid or chromosome), and annotated the correct replicon type of the identified plasmids (Table 1; Methods).

While Table 1 reports the association between a given carbapenemase-carrying plasmid contig and the chromosomal contig with the highest css, the methylation-based css can be calculated between any two contigs to provide the overall contig similarity structure of a metagenomic dataset as the background (Methods). As this can help assess the robustness of associations between plasmid and chromosomal contigs, we visualized the contig similarity structure of each carbapenemase-carrying plasmid contig (M3, M5, and Q1-8; Table 1) in relation to all candidate chromosomal contigs using PCoA (Figure 2; *left*). We also visualised the weighted mean methylation values across the methylation motifs of the five top chromosomal contigs of the top species-classified taxa using heatmaps (Figure 2; *right*). These visualizations strengthen the association predictions with the respective top chromosomal contig shown in Table 1 by showing that other chromosomal contigs of the same taxonomic annotation as the top hit cluster around the plasmid of interest (PCoAs) and by showing the difference in motif methylation ratios between the plasmid and the the top five microbial species (heatmaps).

**Figure 2.**
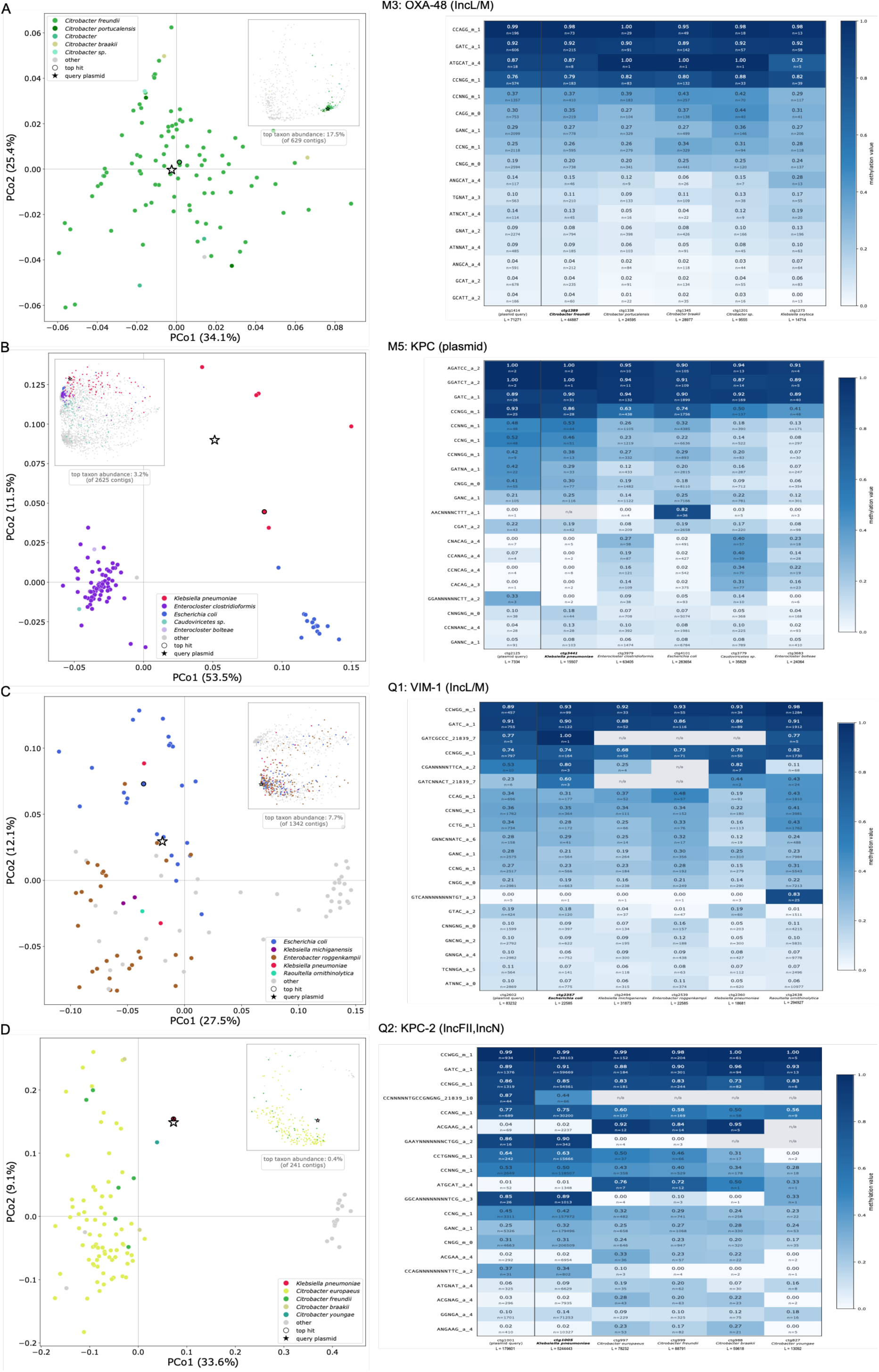
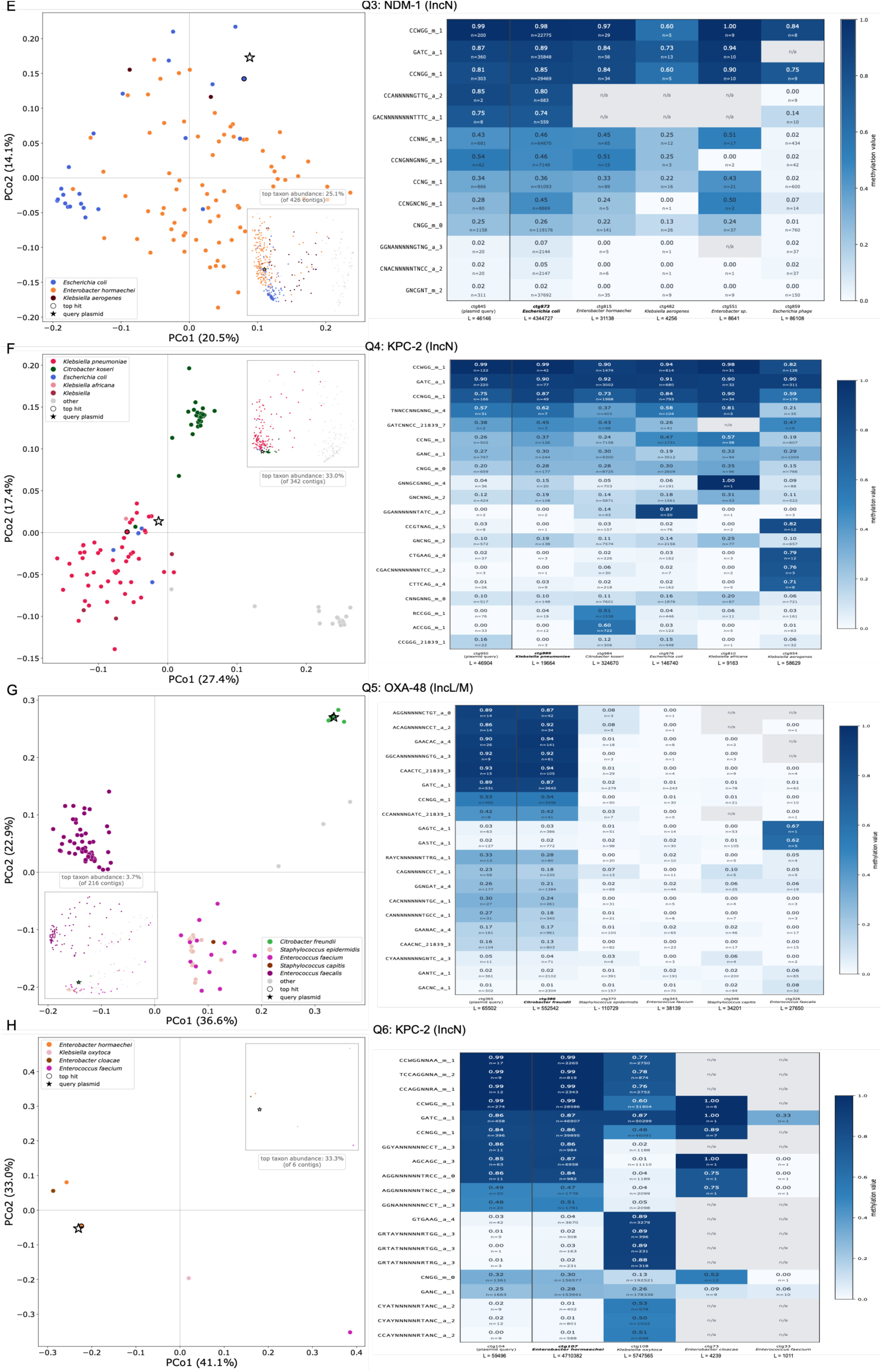
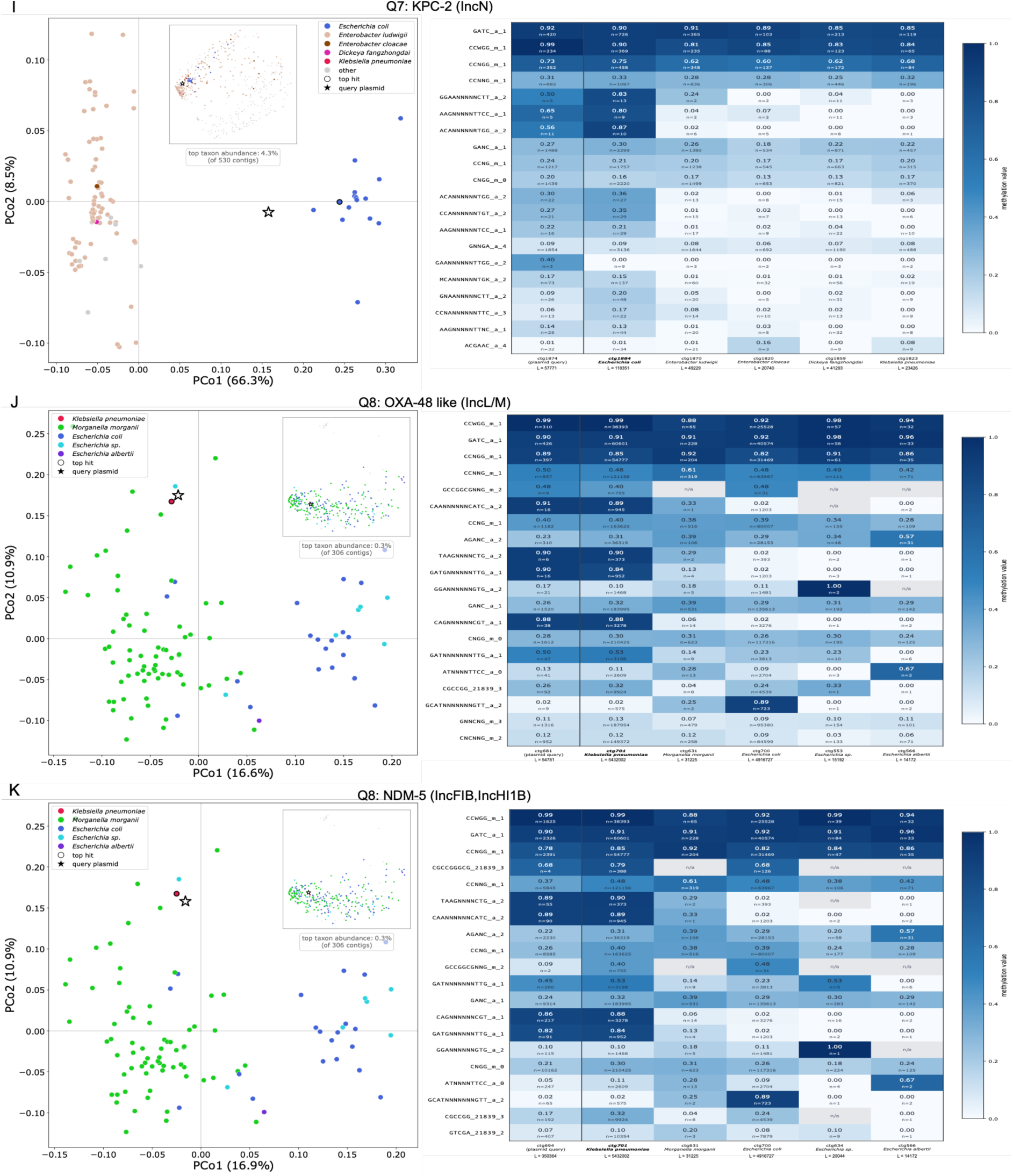
Visualization of methylation-based contig similarity (*left*) and motif methylation (*right*) across the carbapenemase-carrying plasmid contigs (**A-K**) and all chromosomal contigs of their respective dataset. (*left*) PCoAs of the contig similarity structure of each carbapenemase-carrying plasmid contig (star shape; across M3, M5, and Q1-8; Table 1) in relation to all chromosomal contigs. The top 100 chromosomal contigs according to contig similarity score (css) are visualized, respectively; all chromosomal contigs are visualized in the inset. The five top taxa according to css are color-coded, and the top chromosomal contig by css is circled. The relative and absolute top taxon abundance is indicated below the inset. (*right*) Heatmaps of the weighted mean methylation value across up to 20 most-methylated motifs of the respective plasmid contig (leftmost column) and the five top chromosomal contigs of the top species-level taxa. The top species is highlighted in bold.

Some PCoAs reveal that chromosomal contigs of several species cluster around the plasmid contig: For example, in Q1, an *E. coli* chromosomal contig was identified as the top hit (Table 1), but the VIM-1-carrying plasmid contig is also surrounded by *Enterobacter roggenkampii* (part of *Enterobacter cloacae* complex) and *Klebsiella pneumoniae* contigs (Figure 2C), which were both identified as VIM-carriers in the same sample by established diagnostics (Table 1). Similarly, in Q3, an *E. coli* chromosomal contig was identified as the top hit, but the NDM-1-carrying plasmid contig is also surrounded by *Enterobacter hormaechei* (also part of *E. cloacae* complex) contigs (Figure 2E), which was also identified as NDM-carrier in the same sample by established diagnostics (Table 1). Some PCoAs, on the contrary, show that only a single chromosomal contig is the basis of the correct plasmid-microbial host association: In Q6 and Q8, the assemblies recovered only a single chromosomal contig of the correct host (length of 4.71 Mb for Q6, and 5.43 Mb for Q8); the css assigned all three carbapenemase-carrying plasmid contigs of these datasets to their correct chromosomal contigs (Table 1; Figures 2H, J, K). In the case of Q6 (Figure 2H), the assembly led to an overall small number of contigs due to low sequencing depth (Table S1B). In the case of Q8, from which two carbapenemase-carrying plasmids were recovered (Figures 2J, K; *left*), the assembly led to only one contig of the actual host, *K. pneumoniae*., which was correctly identified as the top contig of both plasmid contigs. In such a case of a single correct chromosomal contig, the motif methylation patterns across taxa can further strengthen the association despite the presence of only a single contig as evidence (Figures 2J, K; *right*). In addition, the analysis of sequencing reads in addition to contigs can provide more evidence (see *Application to clinical metagenomics—Read-level associations*).

We additionally annotated all carbapenemase-carrying (quasi)metagenomic plasmid contigs for mobilization potential (Methods). All our plasmid contigs were predicted to be transferable. On these contigs, we identified several additional WGS-confirmed AMR genes (Table S3B), including those conferring resistance to clinically relevant beta-lactams (Table 2). For example, the quasimetagenomic carbapenemase-carrying plasmid contig from Q8 carried all AMR genes that were also detected by the circularized plasmid assembly from the matched isolate’s WGS data (Table S3B). Most plasmid contigs further carried IS elements that flanked the relevant carbapenemases (Table 2).

**Table 2.**
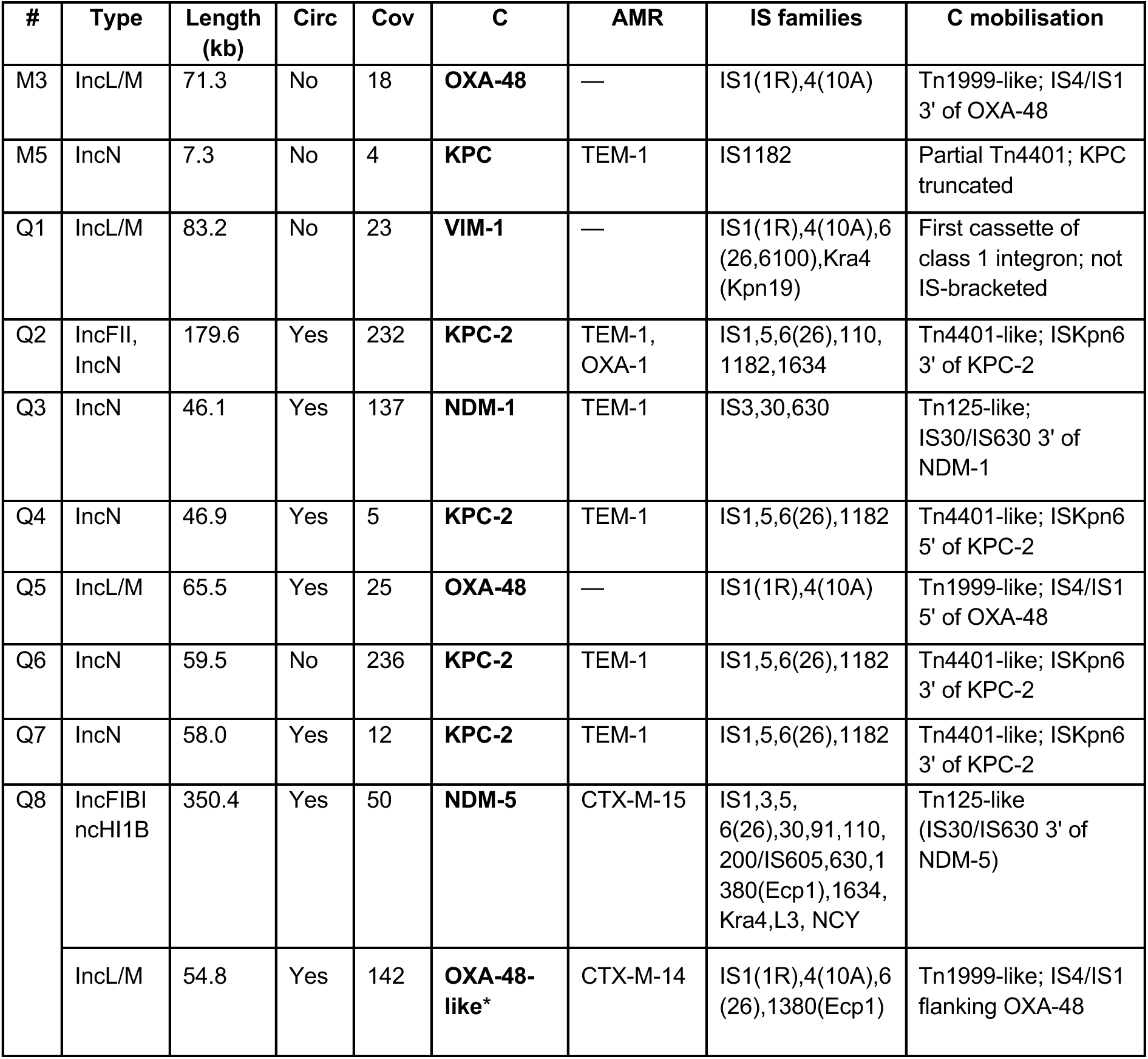
Annotation of plasmid-annotated metagenomic contigs: Replicon type according to MOB-suite annotation; contig length (in kb; see Table S1 for WGS-based plasmid lengths from the same samples), circularity, and mean plasmid contig coverage calculated by nanoMDBG; C as identified carbapenemase gene variant or family; AMR as additionally identified AMR genes using AMRFinderPlus: only genes conferring resistance to beta-lactam antibiotics are reported here (see Table S3B for all AMR genes); IS family identification in a 500b-window upstream and downstream of each C gene; and mobilization context of the C gene. All C-carrying plasmids were further annotated as transferable elements (Methods). *Incomplete sequence recovery precluded allele assignment due to detection at the contig edge (90.9% reference coverage; 100% identical to OXA-48 over the recovered region).

#### Application to clinical metagenomics—Read-level associations

Read-level plasmid-microbial host associations were inferred similarly to contig-level associations (Methods), but as several sequencing reads carried the carbapenemase, we applied a taxonomic majority vote across all sequencing reads (Table S4B). Based on these majority votes, all read-level associations but one were correct (Table 1). In Q4, a single KPC-2-carrying read was associated with *E. coli* while the only *Enterobacterales* grown from this sample was *K. pneumoniae*. Additional two KPC-carrying reads from the same dataset were, however, associated with *K. pneumoniae*, supporting the metagenomic contig- and WGS-based results (Table 1). For all remaining carbapenemase hits, the read-level association results further strengthened the contig-level associations both on the gene variant and the gene family level (Table 1).

In M5, the one KPC-carrying read potentially even recovered the association of KPC with *E. coli*, the second *Enterobacterales* isolate besides *K. pneumoniae* that had been grown from this sample according to established diagnostics and WGS (Table 1); however, the evidence is weak since the KPC gene family could only be found on a single read. Similarly, in Q1, VIM-carrying reads recovered the additional known association between VIM and *K. pneumoniae* besides the contig-based association of VIM with *E. coli* (Table 1; Table S4B). As *K. pneumoniae* also seemed to be a strong candidate on the contig level based on the visualized contig similarity structure (Figure 2C), this provides further evidence for a multi-host association of this particular plasmid—which, in this case, is correct according to our established diagnostics results (Table 1). In addition, read-level analyses enabled the detection of OXA-48 and OXA-48-like genes, which were missed on the contig level, and predicted associations of these genes with two confirmed *Enterobacterales* hosts, *E. coli* (exemplary read: Figure 3A) and *K. pneumoniae* (Figure 3B). In Q6 and Q8, where the assemblies recovered only a single chromosomal contig of the correct host (Table 1; Figures 2H, J, K), the individual sequencing reads provided additional evidence by supporting the association between KPC-2 (n=25)/KPC (n=79) and *E. hormaechei* (Q6; Figure 3C) and between NDM-5 (n=3)/NDM (n=14) as well as OXA-48 (n=16) and *K. pneumoniae* (Q8; Figure 3D).

**Figure 3.**
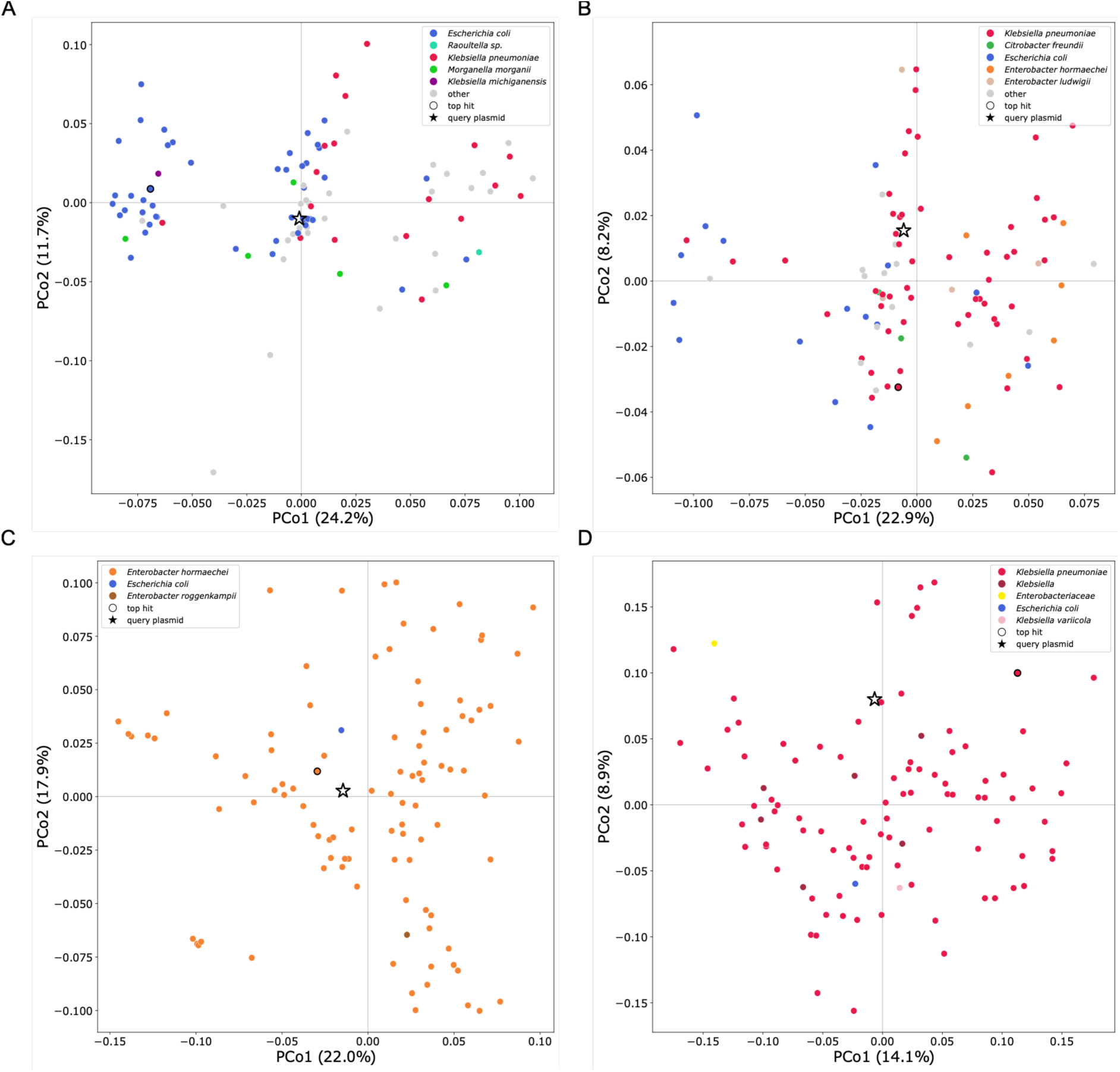
Visualization of methylation-based read similarity of exemplary carbapenemase-carrying plasmid reads: PCoAs of the read similarity structure of a carbapenemase-carrying plasmid read (star shape; Table 1) in relation to the top 100 chromosomal reads according to read similarity score (rss) are visualized, respectively. The five top taxa according to rss are color-coded, and the top chromosomal read according to rss is circled. **A** & **B.** PCoAs of two exemplary OXA-48-carrying plasmid reads in Q1, pointing to associations with *E. coli* (**A**) and *K. pneumoniae* (**B**), respectively. **C** & **D**. PCoAs of two exemplary carbapenemase-carrying reads from Q6 and Q8 that provide robust evidence of plasmid-host associations as a confirmation of contig-level association inferences: (**C**) Association between a KPC-2 carrying read and *E. hormaechei* in Q6; (**D**) Association between an NDM-5 carrying read and *K. pneumoniae* in Q8.

## Discussion

We show that the genomic and epigenetic information inherent to nanopore sequencing data enables the association of clinically relevant carbapenemase genes with their bacterial hosts in metagenomic and quasimetagenomic samples. While the long nanopore sequencing reads facilitate the generation of contiguous assemblies for chromosomal and plasmid identification from complex metagenomic data, methylation signatures encoded in the raw nanopore signal enable the inference of associations between metagenomic assembly-based contigs and individual sequencing reads through similar motif methylation. We leverage this information to attribute carbapenemases as an example of clinically relevant AMR to their pathogenic hosts, first validating our approach using mock communities from clinically relevant bacterial isolates, then applying the optimized analysis pipeline to rectal swab metagenomics and quasimetagenomic data. Comparison with established culture-based diagnostics and isolate assemblies confirms that nanopore metagenomic data can associate both plasmid- and chromosomally encoded resistance genes with their clinically relevant pathogenic hosts in the tested samples.

We provide the first proof that plasmid-host associations derived from clinical metagenomics and quasimetagenomics are feasible and concordant with established diagnostics. This is especially relevant since the only available approach for nanopore-derived methylation-based association is Nanomotif, which works on the metagenomic MAG level to refine metagenomic binning (18,21,22,47). MAG recovery is assembly- and binning-dependent, so low-abundance or divergent taxa can remain unbinned, biasing host–AMR gene links toward abundant organisms (22). Especially in host-dominated clinical metagenomes, pathogen coverage may often be too low for MAG recovery, causing clinically relevant AMR hosts to be missed. This limitation was evident in the rectal swab datasets analysed in this study: Conventional metagenomic assembly and binning recovered a pure MAG of the relevant carbapenem-resistant *Enterobacterales* species in only three of 16 metagenomic or quasimetagenomic samples, and one of these three consisted of a single contig. On the contig and individual sequencing read level, on the other hand, we were able to associate all carbapenemases detected in the metagenomic and quasimetagenomic datasets with their correct bacterial host. We also confirmed these associations by established diagnostics and whole-genome assemblies of the relevant carbapenem-resistant *Enterobacterales* isolates.

The contig-based pathogen detection and AMR association may offer a compromise between bin- and read-level analyses. Contig-level analysis sits between reads and MAGs, retaining AMR gene-carrying sequences that fail binning while providing longer context for taxonomic assignment and resistance gene characterization. It has been shown that metagenomic contigs can be robustly annotated on the bacterial strain level (48), which would enable delineated strain-plasmid associations. We, however, also found that read-level associations of AMR-encoding plasmids and pathogens can provide additional important evidence while only resulting in a single potentially false plasmid-host association in our dataset. Read-level associations can, for example, strengthen weak or unclear contig-level associations (e.g., Q6 and Q6), detect additional genes and associations missed by contig-level associations (e.g., Q1 and M5), and sometimes result in higher gene resolution (e.g., Q8)—depending on where the gene is located on the individual read and how good the assembly quality of the genomic region is. Reads further have the advantage of being biological entities instead of computational constructs such as contigs; contigs might fuse reads of different microbial origins, precluding, for example, the association of one plasmid to several hosts (49). On the read level, evidence can instead be accumulated across reads carrying the same AMR gene; we here show that we can thus predict a potential multi-host scenario of a carbapenemase-carrying plasmid that would have been missed by contig-level analyses but was confirmed by established diagnostics (e.g., Q1).

We further show that metagenomics can provide more granular pathogenic information than established diagnostics. Matched WGS of rectal swab isolates confirmed that metagenomics yields more detailed AMR gene subtype and pathogen strain information than established diagnostics, a finding consistent with previous nanopore WGS studies (8). Crucially, metagenomics can add context unavailable from routine diagnostics: whether an AMR gene is linked to a pathogen, carried on a mobile plasmid, or embedded in a transmissible resistance module. Although this does not yet replace phenotypic AST for therapy, it can guide infection prevention and control escalation, transmission tracing, contact screening, and source investigation (25).

Our study remains limited by several methodological constraints. Validation relied on cultured isolates as the available reference standard, which may miss additional colonizing hosts and underestimate true multi-host plasmid associations. While quasimetagenomic enrichment improved sensitivity in comparison to metagenomics, it may alter community composition and favor organisms that grow rapidly under the enrichment conditions. The contig and read similarity scores are further newly developed metrics, requiring larger benchmarks to define confidence thresholds, failure modes, and reporting standards. Finally, this evaluation focused on carbapenemase-producing *Enterobacterales* from rectal swabs, so performance should be tested across additional sample types, taxa, AMR and other resistance and virulence genes, plasmid backbones, and community structures. Mixed-strain colonization and closely related species complexes might, for example, not allow for bacterial host delineation since strains of the same species may share methylation motifs, assemble into fragmented or composite contigs, or generate ambiguous plasmid-host links. Thus, our approach can support pathogen-host attribution, but strain-level plasmid assignment should be interpreted cautiously unless supported by sufficient strain-specific sequence variation and coverage. This is particularly relevant for clinically important species complexes, such as the *K. pneumoniae* species complex, where routine diagnostics may not reliably distinguish closely related taxa (50). Future benchmarking should include defined same-species and same-complex mixtures, alongside clinical samples with independently confirmed mixed colonization.

In summary, through mock metagenomic communities and application to patient rectal swab data, with cross-validation against established diagnostics and bacterial WGS, we demonstrate that nanopore metagenomics can accurately associate plasmid- and chromosomally encoded AMR genes with their clinically relevant pathogenic hosts. By developing a contig- and read-based approach that overcomes limitations of standard binning, we address an outstanding barrier to metagenomic diagnostics for pathogen and resistance surveillance. Further sensitivity optimizations and diverse benchmarking and prospective clinical validation studies remain warranted to test our approach’s generalizability and establish it for routine surveillance and clinical practice.

## Author contributions

HÜ conducted all computational analyses under LU’s supervision, including data curation and visualization. ES conducted all laboratory work and functional plasmid annotation under LU’s supervision. LU conceptualized the study. HÜ, ES, and LU integrated the computational and clinical interpretation of the data. MB supported the computational interpretation of the data. FG provided samples and laboratory equipment for established diagnostics. EFN, SDB, and RTW consulted on the clinical interpretation. SH and MA consulted on the usage and interpretation of Nanomotif. TR and MJAS contributed to the data analysis. RF contributed invaluable discussions on data interpretation and visualization. HÜ and LU wrote the manuscript with input from all co-authors, especially from ES, RTW, EFN, FM, RS, and RF.

## Ethical approval

Ethical approval for the processing of bacterial isolates and patient rectal swabs was given by the Technical University of Munich ethics committee (2024-522-S-CB). Informed patient consent was waived as samples were taken under routine diagnostics. All human reads from nanopore sequencing were discarded and not analysed beyond quantifying the amount of host DNA generated.

## Acknowledgements

We thank Srinithi Purushothaman for her valuable input on DNA extraction approaches from clinical samples. We thank the laboratory technicians at the Institute of Medical Microbiology at the Technical University of Munich, and especially Hannah Zierer, for their support with sample storage and retrieval and established diagnostics, and the team responsible for the high-performance compute platform at Helmholtz Munich for their support.

## Abbreviations

AMR: antimicrobial resistance
AST: antimicrobial susceptibility testing
BHI: brain heart infusion
BLAST: Basic Local Alignment Search Tool
B: base
CFU: colony-forming unit
Contig: contiguous sequences
Css: contig similarity score
CUPID: Contig- and Unassembled-read-based Pathogen Identification & Delineation
ENA: European Nucleotide Archive
EUCAST: European Committee on Antimicrobial Susceptibility Testing
GPU: graphics processing unit
IS: insertion sequence
Kb: kilobase
MAG: metagenome-assembled genome
MALDI-TOF MS: matrix-assisted laser desorption/ionization time-of-flight mass spectrometry
MLST: multi-locus sequence typing
MPF: mating-pair formation
NCBI: National Center for Biotechnology Information
PBS: phosphate-buffered saline
PCoA: Principal Coordinate Analysis
RMSD: root mean square distance
Rss: read similarity score
ST: sequence type
SUP: super-accuracy basecalling
WGS: whole-genome sequencing

## Supplementary Tables

All supplementary tables are available at https://drive.google.com/drive/folders/1jsUg_eKKBOIJRd8YO7Bb1hX1DYCIslbb?usp=sharing.

**Table S1**. Nanopore sequencing and assembly metrics. **A**. Sequencing, assembly, and plasmid characteristics of the carbapenemase-carrying rectal swab isolates. For each isolate, input DNA concentration, total sequencing yield, mean read Q-score, read count, and read N50 are given. Assembly metrics report the length and median coverage of the chromosomal contig (circularity indicated in parentheses), the detected carbapenemase, and the length, median coverage, and replicon type of the carbapenemase-encoding plasmid contig. Any further plasmid contigs recovered from the isolate are listed under “Additional plasmids” with their replicon type and contig length; “no AMR” denotes plasmids on which no resistance gene was detected. For M5, two isolates were available and are reported as M5a and M5b. For Q8, two carbapenemases (OXA-48 and NDM-5) were carried on separate plasmid contigs, and both are listed. NA: not available. For Q1, no isolate was recovered; therefore, all metrics are reported as NA. **B**. Sequencing and assembly metrics for the metagenomic and quasi-metagenomic rectal swab samples. For each sample, the DNA concentration, input DNA per sample, non-human sequencing yield, mean read Q-score, and Q20 are given, together with read metrics comprising read count, the percentage of reads mapping to the human genome, and the non-human read N50. Assembly metrics report the assembly N50, the number of contigs with the number of circular contigs in parentheses, and the total assembly size. All yield, read, and assembly metrics were calculated after removal of human-mapped reads. M blank and Q blank are the corresponding negative extraction controls. “Too low” indicates a DNA concentration below the reliable quantification limit. NA, not available; metrics not determined for the negative controls. **C**. Isolate sequencing and assembly characteristics and composition of the mock community. The mock community comprised 10 carbapenemase-carrying isolates from defined species and sequence types (STs). For each isolate, the DNA concentration, total sequencing yield, mean read Q-score, read count, and read N50 are given (read metrics), followed by the length and median coverage of the chromosomal contig (circularity indicated in parentheses), the detected carbapenemase, and the type, contig length, and median coverage of the carbapenemase-encoding plasmid contig. Any further plasmid contigs recovered from the isolate are listed under “Additional plasmids” with their contig length and the AMR genes carried; “no AMR” denotes plasmids on which no resistance gene was detected. The final column shows each isolate’s relative abundance in the mock community, calculated as the number of reads for that isolate divided by the total number of reads across all 10 isolates. NA, sequence type not assigned.

**Table S2**. Binning results. Per-bin taxonomic composition of the metagenomic and quasi-metagenomic assemblies. For each dataset, metagenome-assembled bins are listed by bin ID, with the binning tool indicated by the ID prefix (VAMB, MaxBin2, MetaBAT2). For each bin, the total number of binned contigs is given, followed by the absolute taxonomic distribution of those contigs, where each species is shown with its contig count in parentheses and low-count taxa are collapsed into an “other” category. Taxonomic annotation was performed with Kraken2. “Species of interest” denotes the isolate species expected for that sample, and “species of interest %” gives the percentage of contigs within the bin annotated as that species. NA indicates that no contig in the bin was assigned to the isolate species.

**Table S3**. Contig-level plasmid-host associations. **A**. Contig similarity score (css) based on host association of AMR-carrying plasmid contigs in the mock community. For each plasmid contig (ctg ID), the AMR genes carried, the plasmid replicon (Inc) type, and whether the contig was circular are listed; a dash in the Inc-type column denotes no replicon type detected. Host association was performed using the css. The associated host is reported by its taxonomic annotation at two Kraken2 confidence thresholds (0.0 and 0.2); Kraken2 is used only to annotate the css-selected host contig, not to perform the association itself. The number of host contigs sharing the top css is given in parentheses. “Species of origin” gives the true host of the plasmid from the mock community ground truth, and “correct association” indicates whether the css-based assignment matched it. **B**. Css based on host association of AMR-carrying plasmid contigs in the metagenomic and quasi-metagenomic rectal swab datasets. For each plasmid contig (dataset, ctg ID), the AMR genes carried, the plasmid replicon (Inc) type, and whether the contig was circular are listed; a dash in the Inc-type column denotes that no replicon type was detected. Host association was performed using the contig similarity score (css). The associated host is reported by its taxonomic annotation at two Kraken2 confidence thresholds (0.0 and 0.2); Kraken2 is used only to annotate the css-selected host contig, not to perform the association itself. The number of host contigs sharing the top css is given in parentheses. “WGS_confirmed_AMR_genes” lists the resistance genes independently detected by whole-genome sequencing of the corresponding cultured isolate; here, a dash denotes that the isolate did not carry the listed gene(s), and NA denotes a dataset (Q1) for which no isolate was recovered, and confirmation was therefore not possible.

**Table S4**. Read-level plasmid-host associations. **A**. Read similarity scores (rss) in the mock metagenomic community. For each carbapenemase gene family or variant detected on AMR-carrying plasmid reads, the number of carbapenemase-carrying reads (n_reads) and the taxonomic annotation of the chromosomal host read with the highest rss are shown; the number of reads supporting each species-level association is given in parentheses. Species of origin denotes the expected host(s) according to the whole-genome-based ground truth of the mock community isolates, and the final column indicates whether the rss-based majority association recovered the expected host. NDM was carried by two mock-community isolates and was correctly resolved to both *Proteus mirabilis* and *Escherichia coli*. **B**. Rss based on host association of carbapenemase-carrying reads in the metagenomic and quasi-metagenomic rectal swab datasets. For each dataset, only carbapenemase-carrying reads are listed, grouped by the detected carbapenemase gene variant or family. “n_reads” gives the number of carbapenemase-carrying reads, and “top rss” reports the taxonomic annotation of the host read associated by the top read similarity score (rss), with the number of supporting reads for each annotation in parentheses. “WGS-confirmed pathogen” and “WGS-confirmed carbapenemase” denote the pathogen species and carbapenemase, respectively, independently identified by whole-genome sequencing of the corresponding cultured isolate. For Q1, no isolate was recovered; the pathogens and carbapenemase shown for this dataset are instead based on established diagnostics (VITEK2 and MALDI-TOF MS) and are marked with an asterisk.

